# Amelioration of age-related cognitive decline and anxiety in mice by *Centella asiatica* extract varies by sex, dose and mode of administration

**DOI:** 10.1101/2024.01.23.576700

**Authors:** Nora E Gray, Wyatt Hack, Mikah S Brandes, Jonathan A Zweig, Liping Yang, Luke Marney, Jaewoo Choi, Armando Alcazar Magana, Natasha Cerruti, Janis McFerrin, Seiji Koike, Thuan Nguyen, Jacob Raber, Joseph F Quinn, Claudia S Maier, Amala Soumyanath

## Abstract

We have previously reported that a water extract (CAW) of the Ayurvedic plant *Centella asiatica* administered in drinking water can improve cognitive deficits in mouse models of aging and neurodegenerative diseases. Here we compared the effects of CAW administered in drinking water or the diet on cognition, measures of anxiety and depression-like behavior in healthy aged mice.

Three- and eighteen-month-old male and female C57BL6 mice were administered rodent AIN-93M diet containing CAW (0, 0.2, 0.5 or 1% w/w) to provide 0, 200 mg/kg/d, 500 mg/kg/d or 1000 mg/kg/d for a total of 5 weeks. An additional group of eighteen-month-old mice were treated with CAW (10 mg/mL) in their drinking water for a total of five weeks to deliver the same exposure of CAW as the highest dietary dose (1000 mg/kg/d). CAW doses delivered were calculated based on food and water consumption measured in previous experiments. In the fourth and fifth weeks, mice underwent behavioral testing of cognition, anxiety and depression (n=12 of each sex per treatment group in each test).

Aged mice of both sexes showed cognitive deficits relative to young mice while only female aged mice showed increased anxiety compared to the young female mice and no differences in depression were observed between the different ages. CAW (1000 mg/kg/d) in the drinking water improved deficits in aged mice in learning, executive function and recognition memory in both sexes and attenuated the increased measures of anxiety observed in the aged female mice. However, CAW in the diet only improved executive function in aged mice at the highest dose (1000 mg/kg/d) in both sexes and did so less robustly than when given in the water. There were no effects of CAW on depression-like behavior in aged animals regardless of whether it was administered in the diet or the water.

These results suggest that CAW can ameliorate age-related changes in measures of anxiety and cognition and that the mode of administration is important for the effects of CAW on resilience to these age-related changes.

## Introduction

The proportion of people aged 65 and older in the United States (US) was estimated at 16.8% in 2020 and that number is predicted to be over 20%, or about 65 million people, by the year 2040 [1, 2]. By 2030, older people are projected to outnumber children in the US for the first time in history [3]. Aging is associated with a variety of multi-system changes that significantly affect quality of life. Alterations in cognitive function and mood are two such changes that can have a significant impact on the lives of elderly people. A majority of elderly individuals experience some form of memory loss that affects their activities of daily life [4]. These include episodic and source memory deficits [5–7], decreased sensitivity to novelty [8] and impairment in executive function tasks such as attention, planning, inhibitory control and cognitive flexibility [9–11]. Changes in mood are also widespread in the aging population. It is estimated that 29% of people over the age of 55 experience some type of mental health concern, among which anxiety and depression are most common [12]. The proportion of older adults with depressive symptoms increases with age [13] and the prevalence of anxiety disorders is estimated to be as high as 17% in the elderly population. Despite the prevalence of these age-related changes, therapeutic options remain limited, which has led many in the elderly population to seek out alternative therapies. It is estimated that about 60% of older individuals in the US have used at least one complementary and alternative medicine in the last year [14]. In particular, botanical interventions are popular as they have been purported to promote resilience to age-related changes [15, 16].

*Centella asiatica* (CA) is an edible plant used in traditional Chinese and Ayurvedic medicine to improve brain function and boost memory [17, 18] and is marketed in the United States as the dietary supplement “gotu kola” [19]. CA has many reported neurobehavioral effects, including anxiolytic, anti-depressant and cognitive enhancing properties. Anxiolytic and anti-depressant-like effects of the plant have been observed in healthy rodents as well as multiple models of stress including olfactory bulbectomy, chronic immobilization stress and sleep deprivation-induced stress [20–25]. Similar effects have been noted in a handful of human trials where CA treatment reduced anxiety in elderly adults with mild cognitive impairment [26] and decreased symptoms of anxiety and depression in middle-aged adults with generalized anxiety disorder [27]. A substantial literature also supports the cognitive enhancing effects of CA. In human tests, CA extract or dried herb improved cognitive function of healthy middle-aged and elderly [28, 29] individuals as well as elderly subjects with mild cognitive impairment [26]. These effects have been recapitulated in rodent models as well, where CA was shown to improve learning and memory in healthy rats [30] and in mouse models of neurotoxicity [31, 32]. Our own group has also shown that a low dose of the water extract of CA (CAW) in the drinking water can improve cognitive function in mouse models of aging and Alzheimer’s disease [33–36].

In this study, we further investigated the effects of CAW on cognition, measures of anxiety and depressive-like behavior in healthy aged mice. This is the first study evaluating the effects of CAW on anxiety and depression as well as the first to look that effects of increasing concentrations of the extract in aged mice.

In the first part of the study, we evaluated the effects of 5 weeks of treatment with increasing concentrations of CAW integrated in the diet on learning, memory, executive function, measures of anxiety and depressive-like behavior in 3- and 18-month-old C57BL6 mice. We went on to compare the behavioral effects in 18-month-old C57BL/6J mice of the highest concentration of CAW administered in the diet versus in the drinking water.

## Materials and methods

### CAW

CAW was prepared as previously described by our group [37]. Briefly, CA (dried aerial parts; 4 kg) purchased through Oregon’s Wild Harvest (Redmond, OR, USA) was boiled with deionized water (50 L) for 90 min. The mixture was allowed to cool until safe to handle, allowing plant material to settle to the bottom of the boiling kettle. The upper liquid was filtered to remove insoluble materials including finer plant debris (McMaster-Carr #5162K112 filter bag). The filtrate was frozen in aluminum baking trays and lyophilized in 3 separate batches to yield extracts BEN–CAW-7, 8 and 9 (total weight of 820 g) all of which were used during the course of the present study. Voucher samples of the starting plant material are stored at the Oregon State University Herbarium (Voucher number OSC-V265416) and in our laboratory at Oregon Health & Science University (Voucher number BEN-CA-6), along with voucher samples of the dried CAW extract batches (Voucher numbers BEN-CAW 7, 8 and 9).

### Animals

C57BL6 mice were obtained from the National Institute of Aging Aged Rodent Colony. Mice were kept in a climate-controlled environment with a 12-h light/12-h dark cycle. Water and diet (AIN-93M; Dyets Inc., Bethelem, PA, US) were supplied *ad libitum*, except during the Odor Discrimination Reversal Learning testing when food was restricted at night and resupplied in the afternoon following testing. All methods were performed in correspondence with the NIH guidelines for the Care and Use of Laboratory Animals and were approved by the Institutional Animal Care and Use Committee of the Portland VA Medical Center.

#### CAW in diet and drinking water

Diets containing CAW were prepared by Dyets Inc. by mixing CAW (at 0.2%, 0.5% and 1.0 % w/w) with the AIN-93M diet components until a homogenous distribution was achieved. After adding cold water (10%), the diet was run through a California Pellet Mill CL-3 lab pellet mill to create pellets, which were air dried at 27°C for 24 hours. Finally, the diet was sterilized by gamma irradiation (5.0-20.0 kGy; Sterigenics, Oak Brook, IL, USA).

For administration in drinking water, CAW was dissolved in deionized water at a concentration of 10 mg/mL. Drinking water containing CAW was replaced twice weekly, *i.e.*, every 3 or 4 days.

#### Composition and content of CAW and CAW-containing diets

The chemical composition of the CAW used was previously reported [37]. Inclusion of CAW in the rodent diets at the required levels was verified using liquid chromatography multiple reaction monitoring mass spectrometry (LC-MRM-MS) of triterpene and caffeoylquinic acid components as previously reported by our group [37]. We also verified that there was no microbial contamination and confirmed the stability of those compounds in the diet over several months storage to ensure that degradation was not occurring (Supplementary Tables 1 and 2).

### CAW Treatment

The treatment paradigm is outlined in Figure 1. Briefly, three- and eighteen-month-old mice were kept on AIN-93M treated with CAW for a total of 5 weeks. Young (3-month-old) and old (18-month-old) mice were treated with CAW integrated into AIN-93M at concentrations calculated to deliver 0 mg/kg/day, 200 mg/kg/day, 500 mg/kg/day and 1000 mg/kg day exposure based on average mouse weights and food consumption from previous studies [38, 39]. An additional treatment group of old mice exposed to CAW in the drinking water (10 mg/mL) for 5 weeks was also included. This concentration of CAW in the water was selected to approximate the 1000 mg/kg/day exposure again based on calculations from previous studies [38, 39]. The number of animals in each treatment group can be found in Table 1. The duration of the behavioral test required splitting the mice into two cohorts so that testing for each mouse did not exceed two weeks. To accomplish that each treatment group was divided in half so that 12 of each sex were evaluated for each behavioral endpoint described below.\

**Figure 1:**
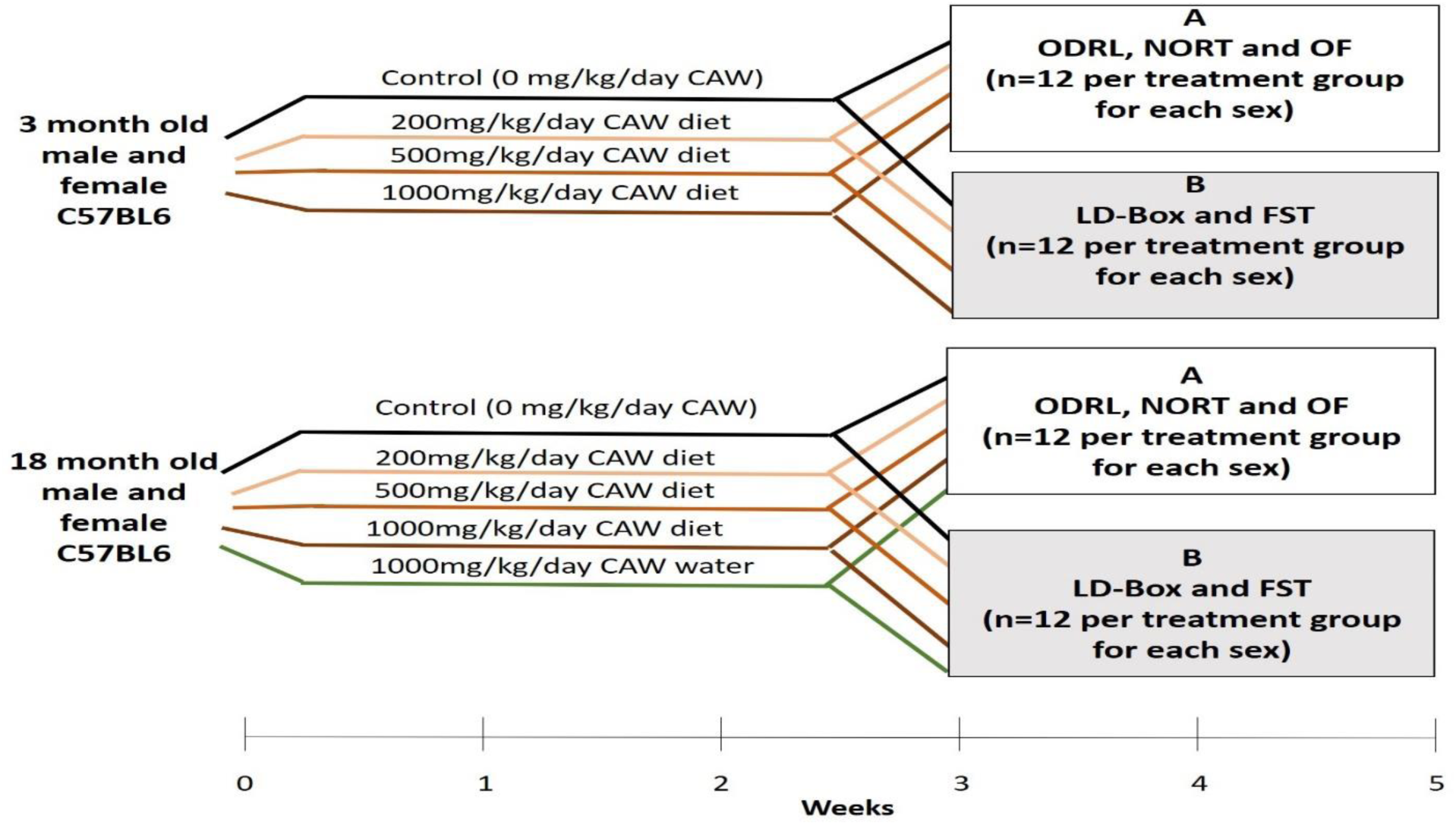
Treatment timeline and tests applied to each of two subgroups groups of mice receiving similar treatments. Mice in group A underwent ODRL: odor discrimination reversal learning, NORT: novel object recognition test and OF: open field. Mice in group B underwent LD-Box: light dark box and FST: forced swim test Group A: odor discrimination reversal learning (ODRL), novel object recognition test (NORT) and open field (OF) or Group B: light dark box (LD-BOX) and forced swim test (FST).

**Table 1:** Number of animals in each treatment condition.

| Age of mice | 0 mg/kg/day<br>CAW | 200<br>mg/kg/day<br>CAW in the<br>diet | 500<br>mg/kg/day<br>CAW in the<br>diet | 1000<br>mg/kg/day<br>CAW in the<br>diet | 1000 mg/kg/day<br>CAW in the<br>drinking water |
| --- | --- | --- | --- | --- | --- |
|  | Male/Female | Male/Female | Male/Female | Male/Female | Male/Female |
| 3 month old | 24/24 | 24/24 | 24/24 | 24/24 | 0/0 |
| 18 month old | 24/24 | 24/24 | 24/24 | 24/24 | 24/24 |

### Behavioral testing

In the final two weeks of treatment, mice underwent behavioral testing. Mice (n=12 per treatment) were subjected to one of two groups of behavioral tests (Figure 1),

Although 12 male and 12 female receiving each treatment condition were a for allocated for a given test, data was not collected for that many animals in every behavioral test due to a variety of factors, including technical issues with the software, non-participation of mice in the task, and COVID-related issues with the experimenters. The number of animals that completed each test can be found in Supplementary Table 3.

*ODRL:* This test assesses learning and cognitive flexibility [40]. It consists of three separate stages: shaping, acquisition and shift. In the shaping stage, mice were introduced to the testing chamber and trained to dig for a food reward in lavender scented bedding. Mice were presented with one cup containing the food reward that had been successively filled with bedding in five different levels; 0%, 25%, 50%, 75% and 100%. The mouse proceeded to the acquisition step when it successfully retrieved the reward five times in succession.

The acquisition stage followed the completion of the shaping stage. Mice were presented with two cups, one that contains dried beans and one with string. In every trial, one digging material was scented with a mint odor and the other with vanilla. The odor and material pairings were randomly alternated between trials but balanced throughout acquisition phase so that each mouse was exposed to roughly equal combinations of each odor and digging material. The position of the baited cup, whether it was presented on the right or left side of the apparatus, was also balanced throughout testing. During the acquisition stage the bowl with the mint odor was always baited, regardless of the digging material. Example trials are outlined in Table 2. The criteria for completing the acquisition phase was 8 correct digs in any bout of 10 trials. The number of trials for each mouse to reach criteria was recorded.

**Table 2:** Example of test pairings for Odor Discrimination Reversal Learning (ODRL) test: Representative combinations of odor and digging material pairings during each phase of the ODRL. D1: dried beans; D2: pieces of string; O1: vanilla; O2 = mint. Italicized and bold indicates the baited cup.

|  | Right position | Left position |
| --- | --- | --- |
| Acquisition phase | D1 + O1 | **D2 + O2** |
|  | **D1 + O2** | D2 + O1 |
|  | D2 + O1 | **D1 + O2** |
|  | **D2 + O2** | D1 + O1 |
| Shift phase | **D1 + O1** | D2 + O2 |
|  | **D1 + O2** | D2 + O1 |
|  | D2 + O1 | **D1 + O2** |
|  | D2 + O2 | **D1 + O1** |

After a mouse reached the criteria in the acquisition phase, it immediately proceeded to the shift phase. As in the acquisition phase, in the shift phase, mice were presented with two cups, one containing dried beans and the other string. As before, in each trial, one digging material had the vanilla odor and the other the mint odor, and again the odor + digging material pairings were balanced throughout the trial as was right/left location of the baited cup. In the shift phase, however, the cup with the dried beans was always baited regardless of odor so that the mouse had to ignore the previously learned association with odor and now learn to associate the food reward with the digging material. Again, criteria was defined as 8 correct trials in any bout of 10 and trials to criteria was recorded.

Mice were food restricted the night before each phase of the ODRL to motivate the animals. (what is important here is whether the rewarded condition involved novelty or not as we are trying to compare this to the object recognition test involving novelty; mice anxious to explore novelty might lead to reduced exploration of the novel object and increased exploration of the familiar object. Looking at the data, we clearly see that some mice very much prefer the familiar object; so it is not that these mice do not distinguish the objects but they prefer the known/familiar one. This aspect of novelty might be less or not playing a role in the ODRL and that might explain the difference; would recommend incorporating something along these lines in the discussion)

*NORT:* This is a test of recognition memory [41] that also has three stages: habituation, training and testing. In the habituation phase, each mouse was introduced to an empty square arena (38 x 38 x 64 cm, built of white acrylonitrile butadiene styrene) for two 10-minute sessions on two consecutive days. During the subsequent training stage, animal are exposed to two identical objects for 10 minutes, once an hour in a 3 hour period. The testing phase occurred 2 hours and 24 hours after the final training phase. In the testing phase, one of the identical objects was replaced with a novel object and the mouse was given the chance to examine the environment for 5 minutes. The time spent exploring the familiar and novel objects was evaluated via a camera placed above the arena, interfaced with ANYmaze video tracking system (Stoelting Co, Wood Dale, IL). At 24h, mice were reintroduced to the arena again and presented with the same familiar object from training and the first test session along with a second novel object (distinct from the object used in the 2h test). Again, the time spent exploring the familiar and novel objects was evaluated.

*OF:* The OF test can be used to assess anxiety and overall activity [42]. In the OF test, each mouse was placed into a square arena (38 x 38 x 64 cm high, constructed of white acrylonitrile butadiene styrene (ABS)) for a 10-minute open field session. A camera mounted above the arena, interfaced with a video tracking system (Any-maze, Erie, PA), captured time immobile (s) and time in the center (s). Increased time in the center and reduced time immobile are associated with decreased measures of anxiety.

*LD-Box:* The testing chamber consists of a square Plexiglas chamber (60cm x 60cm x 26cm) fitted with infrared beams on all sides to measure locomotor activity and a z-axis beam to measure rearing. This test is intended to test for measures of anxiety by placing a dark insert into the chamber (16”x16”x12”). Additional lighting is provided on the open portion to achieve a lux of 3000 and a cutout door (4”x2”) allows the animal to freely move between zones. Mice are placed in the light portion of the light/dark box and time in each zone, overall activity, and latency to enter the dark were measured with the infrared beams for each zone of the apparatus (light or dark). The test lasted for a total of ten minutes. Increased time in the light portion of the arena reflects decreased measures of anxiety [43].

*FST:* The forced swim test assesses depressive-like behavior [44]. The forced swim test occurred in a plastic tub (22 cm H x 28 cm W x 40 cm L) filled with lukewarm tap water at 25 °C [45]. The test occurs over two days with the time spent immobile over a 5-minute period measured each day. Results reported are from the second day of the test when immobility is more pronounced than on the first day. Increased immobility is interpreted as increased depressive-like behavior.

*Statistical analyses:* Prior to formal analysis, we visually examined the outcomes to check for distribution patterns and possible outliers. No outliers were removed in our analysis. Bivariate plots were created to investigate how the endpoints relate to grouping variables. We then built a linear regression model with a three-way interaction involving treatment, age, and sex. Likelihood ratio tests were conducted to determine the significance of interactions related to sex. If significant, we stratified the analysis by sex and ran a two-way interaction model for treatment and age. Otherwise, we included sex as a covariate. If linear regression wasn’t suitable due to issues like heteroskedasticity and non-normal residuals, we used a generalized linear model (GLM) following the same approach. For endpoints involving percentages or discrimination indices, we transformed the data into proportions and applied a beta model. Otherwise, we utilized a GLM with a log-link and Eicker-Huber-White standard errors. A summary of which analyses were conducted for which endpoint can be found in Supplementary tables 4 and 5.

We conducted *a priori* contrasts and applied an MVT adjustment to control family-wise error rates when comparing p-values. All analyses were conducted using the betareg [46] and emmeans [47] packages in R V4.3.0.

For endpoints with a significant interaction between age, treatment effect and sex, male and female mice were graphed separately. For endpoints without significant interactions, both sexes were plotted together. All graphs show individual data points and means and error bars reflect the standard error of the mean.

## Results

### CAW administered in the diet improves learning and cognitive flexibility in aged mice in a dose-dependent manner

The ODRL is a test of learning and the cognitive flexibility domain of executive function. An increased number of trials to reach criteria in the acquisition phase suggests deficits in learning, while the same increase in the shift phase indicates impairments in cognitive flexibility. There was no significant interaction between age, sex and treatment in either the acquisition or shift phase of the ODRL and so male and female mice were analyzed together. Eighteen-month-old mice exhibited worse performance, compared to three-month-old mice, in both the acquisition (Figure 2A) and shift (Figure 2B) phases of the ODRL. Treatment with CAW integrated into the diet improved performance in both phases for old, 18-month-old, animals and the magnitude of improvement increased with increasing concentrations of CAW (Figure 2A and 2B).

**Figure 2:**
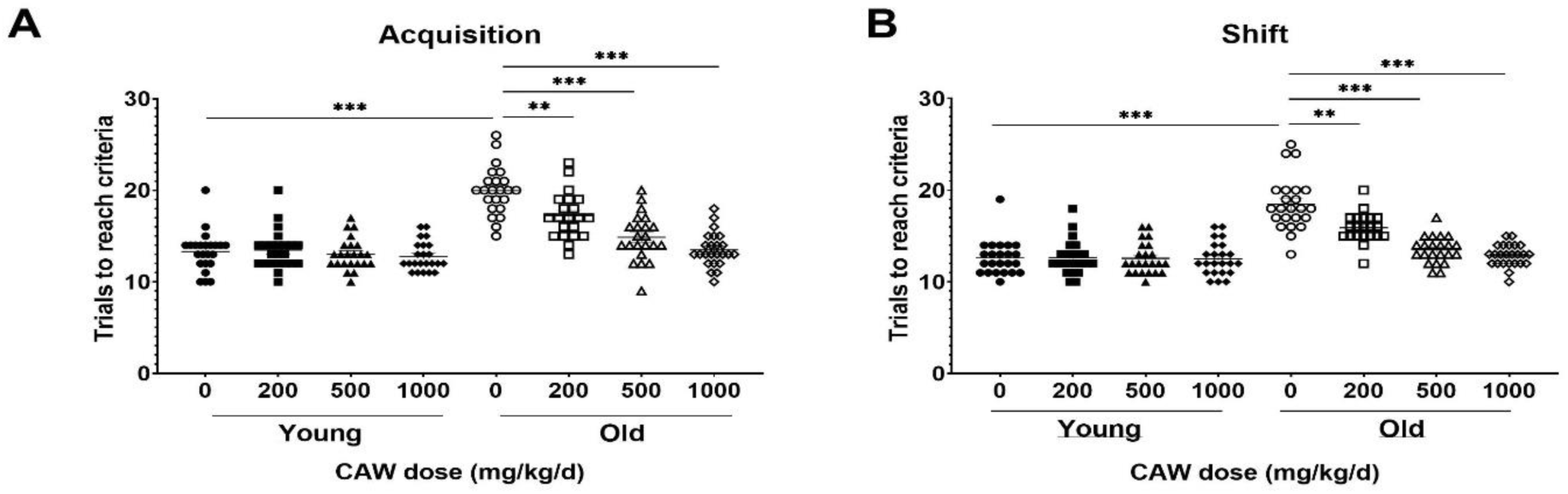
CAW in the diet improves learning and executive function in old mice in a dose-dependent manner. Deficits were seen in the acquisition (A) and shift (B) phases of the ODRL in old mice relative to young animals, which were attenuated with CAW treatment administered in the diet. Increasing concentrations of CAW resulted in greater improvements in ODRL performance. Combined data from male and female mice are shown as sex differences were not observed. **p<0.01, ***p<0.0001

In contrast, CAW treatment administered in the diet had no effect on the performance of young, 3-month-old, mice in either the acquisition or shift phases of the ODRL regardless of concentration (Figure 2A and 2B).

### CAW in the diet has divergent effects on recognition memory in male and female old mice

The NORT test of recognition records the amount of time the mice spent exploring a novel object versus a familiar one. Decreased time spent with the novel object indicates poorer recognition memory. There was a significant interaction between sex, age and treatment in both the 2- and 24-hour retention tests. At the 2-hour time point, no significant differences were seen for female mice in the percent time spent with the novel object regardless of age or treatment, although in young female mice there was a non-significant trend towards reduced time with the novel object seen in animals treated with 500 mg/kg/d (Figure 3A). At the same time point in male mice, there was a significant improvement time spent with the novel object in aged mice treated with 500 mg/kg/d CAW, an effect not observed in young mice (Figure 3B). In contrast, in aged female mice 500 mg/kg/d CAW improved NORT performance at 24 hours (Figure 3C). In male mice this concentration of CAW had no effect, although 200 mg/kg/d CAW actually decreased time spent with the novel object in aged male mice (Figure 3D). CAW treatment in the diet did not impact performance in young animals of either sex regardless of concentration.

**Figure 3:**
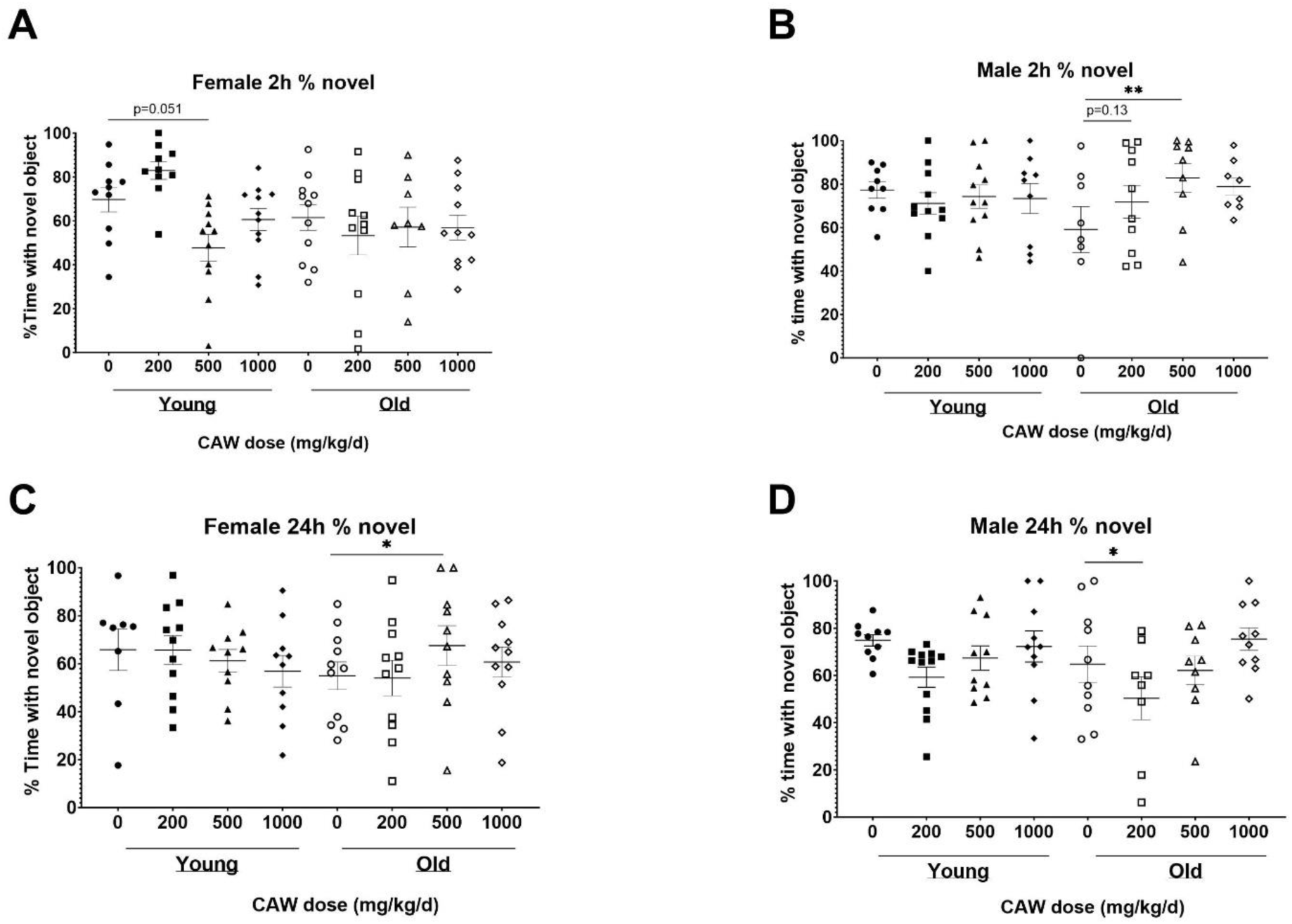
CAW in the diet results in inconsistent modulation of NORT performance. There was a significant interaction between sex, age and treatment in the NORT. A) No significant effects of age or treatment were observed in female mice during the 2h test. B) In aged male mice, 500 mg/kg/d CAW improved performance during the 2h test. At 24h, C) 500 mg/kg/d improved NORT performance in aged female mice while D) 200 mg/kg/d resulted in impaired performance in aged male mice. *p<0.05, **p<0.01

### CAW administered in the diet has no effect on behaviors associated with measures of anxiety and depressive-like behavior in old mice and may increase some anxiety-like behavior in young female mice

The OF test of anxiety monitors total time spent in the center of the field as well as time immobile. Decreased time in the center reflects enhanced anxiety. Increased time immobile can also reflect increased anxiety although that effect depends on where in the field (center vs periphery) the immobility occurs. There was no interaction between age, sex and treatment in time immobile in the OF. No differences in time immobile were observed between young and old control mice nor did CAW treatment affect the time immobile at any concentration at which it was administered in the diet in either the old or the young mice (Supplementary Figure 1A).

There was a significant interaction between age, sex and treatment effect for the time in the center in OF. Aged, control female mice displayed significantly decreased time in the center compared to young, female, control animals, although CAW treatment did not affect this endpoint in the old female mice at any concentration (Figure 4A). Interestingly, all concentrations of CAW significantly reduced time in the center in young female animals (Figure 4A). In contrast, in male mice no effects of either age or treatment were evident in the time in the center in the OF (Figure 4B)

**Figure 4:**
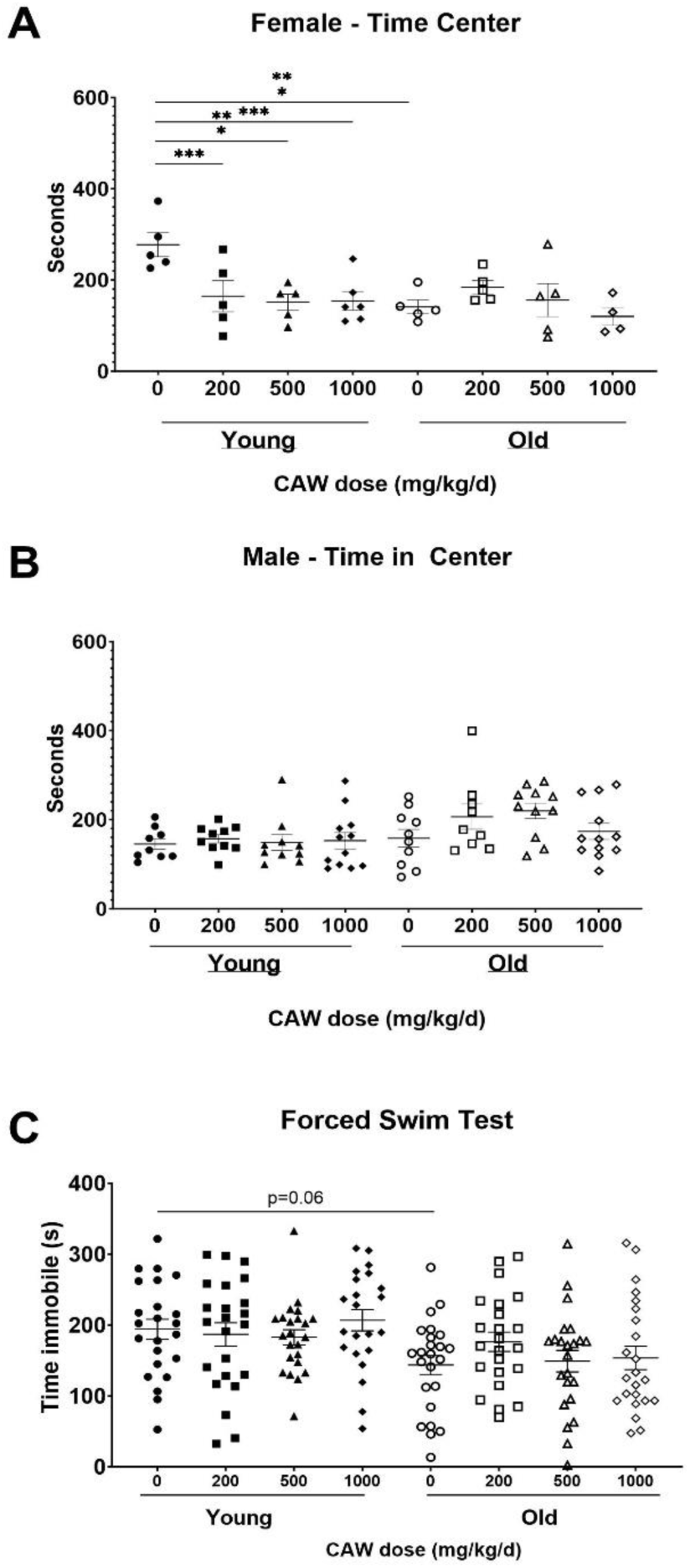
CAW in the diet does not improve measures of anxiety and depression. There was a significant interaction between sex, age and treatment in the time in the center for the Open Field (OF) test.A) Aged female mice displayed reduced time in the center relative to young mice. CAW did not affect time in the center in old mice at any concentration but higher concentrations of CAW resulted in reduced time in the center of young female mice. B) There were no effects of age or treatment on time in the center for male mice. There were similarly no effects of age or treatment on time immobile in the FST (C). ***p<0.001

LD-Box is another test that evaluates anxiety. More anxious mice will spend more time in the dark than less anxious mice. In this test, however, we did not observe any differences between groups regardless of age or treatment (Supplementary Figure 1B).

The FST evaluates depressive behavior. Increased time immobile indicates increased depressive behavior. Here again no significant effects of age, sex nor of any concentration of CAW administered in the diet were detected although surprisingly there was a non-significant trend towards reduced time immobile in the old control mice relative to the young control animals (Figure 4C).

### CAW given in the drinking water elicits even greater attenuation of age-related impairments in learning and executive function in aged mice than CAW given in the diet

We assessed the effects of 5 weeks of treatment with the highest dose of CAW (1000 mg/kg/d equivalent) administered in the diet compared with when it was administered in the drinking water. In the acquisition phase of the ODRL, we found that CAW in the drinking water attenuated age-related deficits even more robustly (p<0.05) than when it was given in the diet (Figure 5A). The same enhanced response in old mice to CAW in the drinking water compared to the diet (p<0.05) was observed in the shift phase of the ODRL (Figure 5B).

**Figure 5:**
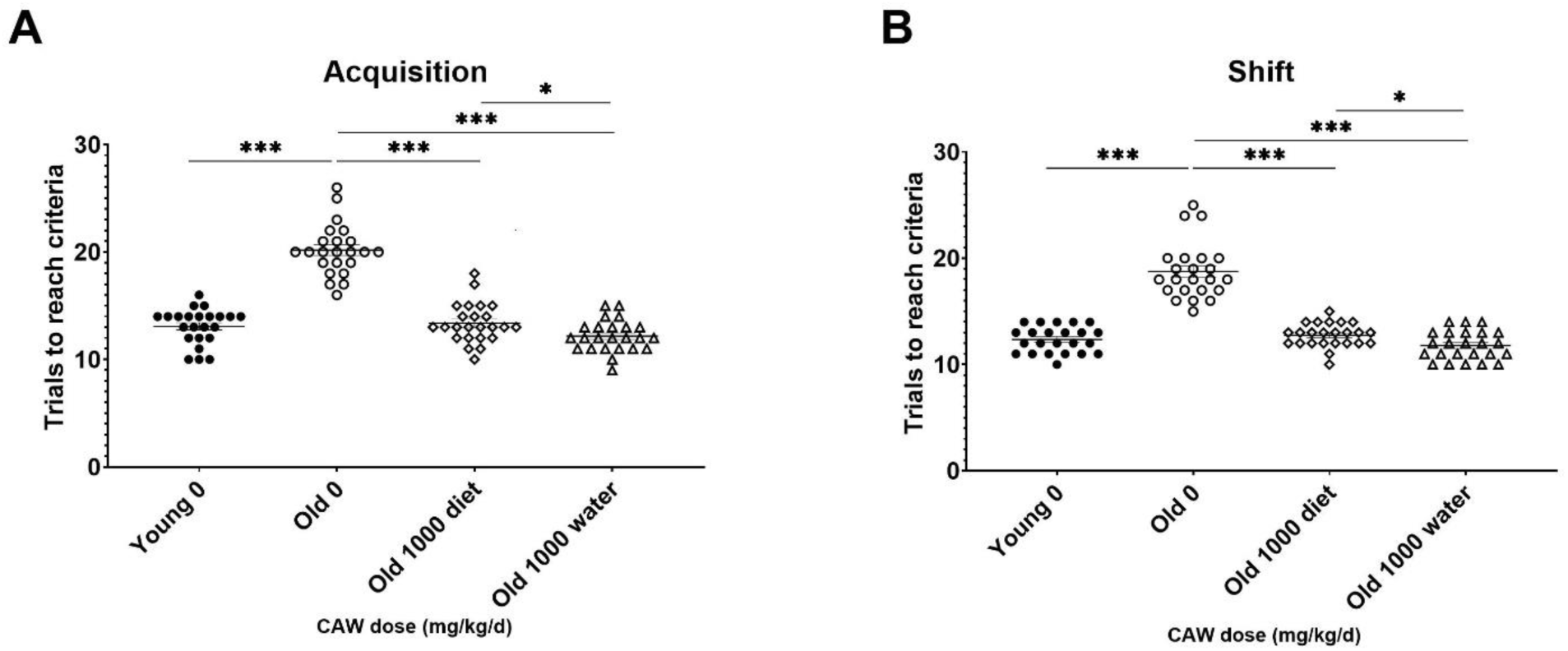
CAW in the drinking water improves learning and executive function in old mice more robustly than when given in the diet. CAW attenuated age-related deficits in ODRL performance in both the acquisition (A) and shift (B) phases of the ODRL. Improvements elicited by CAW in the drinking water were even greater than those seen with CAW treatment in the diet. *p<0.05, ***p<0.0001

### CAW in the drinking water improves recognition memory in aged animals

When CAW was administered in the drinking water, there was a significant increase in the percent time spent with the novel object apparent in aged mice during the two-hour retention test (Figure 6A). A similar but non-significant was observed in aged mice receiving CAW in the drinking water at 24h (Figure 6B).

**Figure 6:**
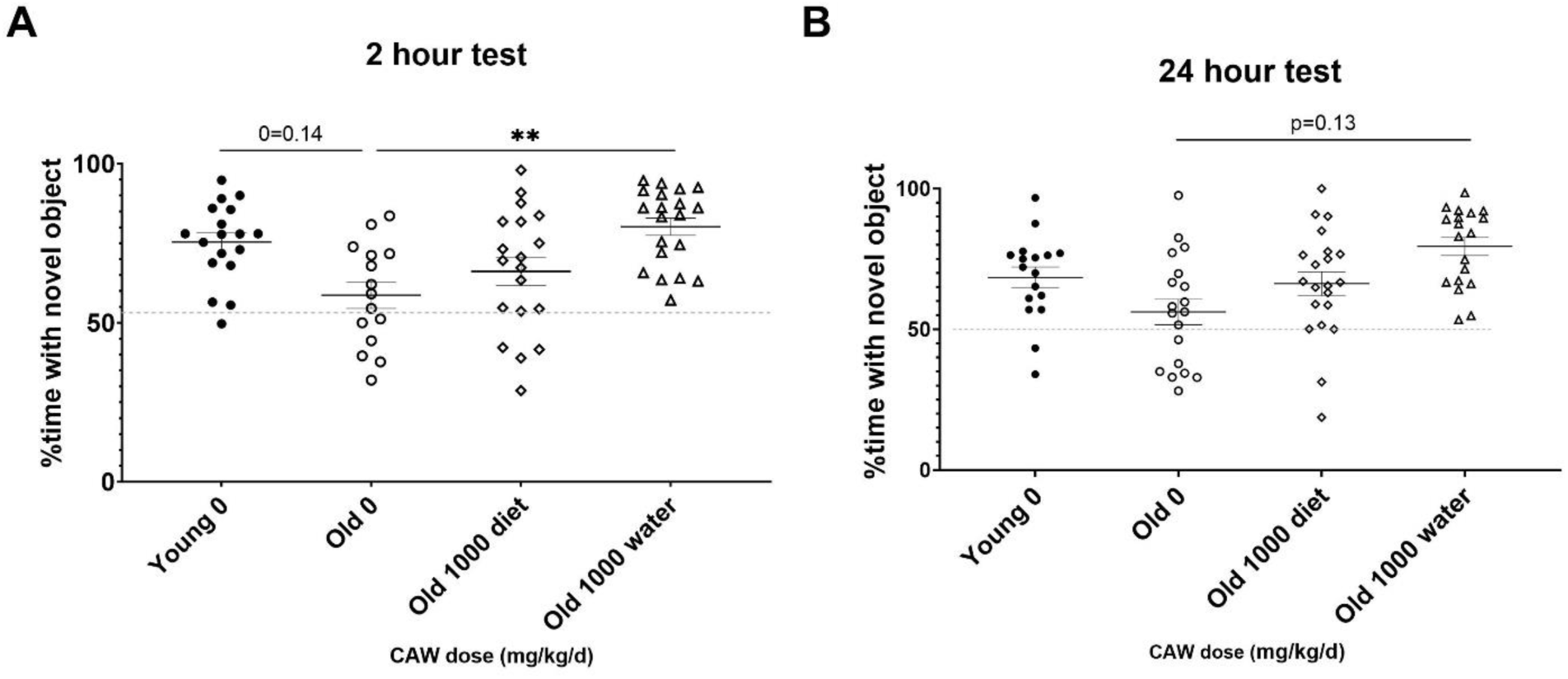
Recognition memory in aged mice is improved by CAW treatment in the drinking water but not in the diet. CAW given in the drinking water (A) significantly increased time spent with the novel object at 2h and (B) showed a trend to an increase at 24h. **p<0.01

### CAW given in the drinking water has inconsistent effects on different metrics of anxiety but does not affect depressive behavior in aged mice

There was a significant interaction between age, sex and treatment effect when examining time in the center of the OF test. There was a robust increase in time in the center in aged female mice in response to CAW given in the drinking water compared to aged female mice that either received CAW in the diet or did not receive any CAW (Figure 7A). This suggests a decrease in age-related anxiety in the female mice.

**Figure 7:**
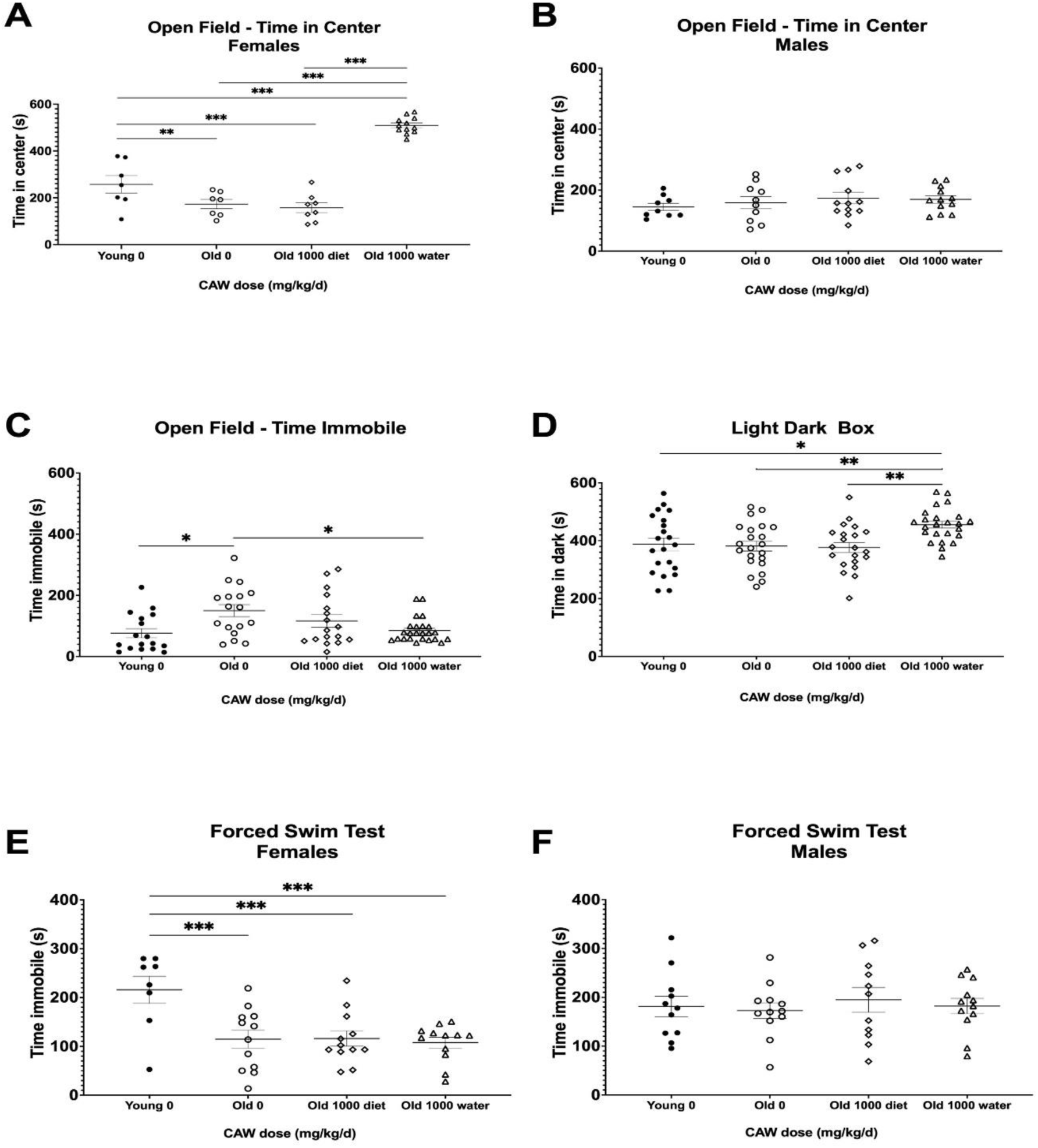
Although CAW in drinking water improves some metrics of anxiety in aged mice, it exacerbates other metrics and has no effect on depression-like behavior. There was a significant interaction between sex, age and treatment in the time in the center for the OF. A) Aged female mice displayed reduced time in the center relative to young mice. CAW in the diet did not affect time in the center in old mice but CAW in the drinking water resulted in increased time in the center for old female mice. B) There were no effects of age or treatment on time in the center for male mice. C) CAW in the drinking water attenuated increases in time immobile in the open field in old mice while CAW in the diet did not significantly affect this endpoint. D) In aged mice CAW the diet did not affect time in the dark in the LD-Box but CAW in the drinking water actually increased time in the dark. D) In the FST, aged female mice displayed reduced time immobile relative to young female mice. CAW treatment had no effect on time immobile in the FST whether administered in the drinking water or the diet. *p<0.05; **p<0.01; ***p<0.001.

However, there was no effect of CAW in the drinking water on time in the center for male mice (Figure 7B). There was no interaction between age, sex and treatment effect for time immobile in the OF. CAW in the drinking water significantly attenuated the age-related increase in time immobile in aged mice in the OF test, while CAW in the diet did not affect this endpoint (Figure 7C).

In contrast, when using the LD-Box to assess anxiety, we observed an increase in time in the dark in aged mice following CAW administration in the drinking water (Figure 7D). In this test however age-related changes were not observed.

In the FST test, there was an interaction between sex, age and treatment for time immobile. Aged female mice had reduced time immobile relative to younger mice on day 2 (Figure 7E) but the same age effect was not seen in male mice (Figure 7F). CAW had no effect on FST time immobile in either sex regardless of the route of administration (Figure 7E and 7F).

## Discussion

Alterations in cognition and mood are highly prevalent in the aging population and negatively affect quality of life. Approximately two out of three Americans experience at least some form of cognitive impairment by age 70 [48]. Changes in mood are similarly common in the aging population, with a prevalence of depressive symptoms and anxiety of nearly 1 in 5 in elderly people [12, 13]. All of these conditions also often severely curtail daily life activities in older people further underscoring the need for therapies that can increase resilience to these age-related challenges.

The cognitive, antidepressant and anxiolytic effects of *Centella asiatica* have been demonstrated many times in both rodent and human studies [20–24, 26–29, 32]. Our group has previously shown that the water extract of *Centella asiatica* (CAW) administered in the drinking water at 2 mg/mL (calculated to deliver 200 mg/kg/d CAW) improves cognitive function in mouse models of aging and neurodegenerative disease [33–36, 38]. This study builds on that previous research to explore the ability of CAW to reverse age-related changes in anxiety and depression. We also evaluated the effects of multiple doses of CAW on cognitive and mood endpoints and assessed sex differences in response to the extract.

In our study, we observed impaired performance in aged mice (compared to young mice) in ODRL, NORT and OF, although for OF the age effect was only observed in female mice. The impaired performance we observed in these tasks is in line with previous reports that also demonstrated diminished ODRL, NORT and OF performance in aged mice relative to young ones [49–51]. We did not, however, observe any age-related deficits in LD-Box or FST or for male mice in the OF. This is in contrast to previous reports in the literature where poorer performance was observed in LD-BOX, FST and OF [50, 52, 53] and could be related to differences in the ages tested or it might reflect subtle differences in testing protocol and metrics recorded. For instance, in one study using LD-BOX, reduced distance traveled in the dark was seen in 23-month-old mice but not 17-month-old mice as compared to 3-month-old animals [50]. In the same study time immobile in the FST test was increased in 24-month-old mice relative to 3-month-old animals but no differences were seen between the 3-and 18-month-old groups [50]. This suggests that perhaps a larger difference in age is required to observe differences in performance in the FST and LD-BOX.

When CAW was administered in the diet, we observed dose-dependent attenuation of age-related deficits in learning and cognitive flexibility. Old mice receiving the highest concentration of CAW in the diet (1000 mg/kg/d) showed improved performance in the ODRL test compared to what was seen in the young, 3-month-old mice. The effect of CAW in the diet on recognition memory was less robust. Significant improvements were seen in old male mice treated with 500 mg/kg/d in the 2h NORT test and in old female mice treated with the same concentration in the 24h test but no significant deficits were detected between old controls and young controls. Surprisingly, there was also a significant impairment in NORT performance in the old male mice treated with 200 mg/kg/d in the 24h test. The lack of a consistent effect of CAW in the diet on NORT performance is in contrast to the clear benefit observed in the ODRL. It is possible this discrepancy could be explained by a differential effect of CAW in distinct brain regions. The ODRL is mediated by the medial prefrontal cortex [54, 55], while NORT performance relies on inputs from perirhinal cortex, hippocampus, medial prefrontal cortex and medial dorsal thalamus [56]. While further studies are needed to confirm that the levels of CAW constituent compounds are consistent across various brain regions, we feel that differences are unlikely possibility given our previous report of similar effects of CAW on antioxidant gene expression in the cortex, hippocampus and cerebellum of aged mice [33]. It is more probable that the different effects seen in the ODRL and NORT reflect differences in the sensitivity and variability of each test. We recognize that anxiety towards exploring novel objects (neophobia) might have contributed to these divergent findings. We observed a much greater amount of variability between animals in NORT performance which likely prevented the detection of more significant and consistent findings.

CAW given in the diet did not appear to affect metrics of anxiety in aged mice. However, there was a statistically significant decrease in time spent in the center of the open field in young female mice at all concentrations of CAW tested, suggesting elevated levels of anxiety. This effect in young mice and the lack of effect in aged mice was surprising given the purported anxiolytic effects of CAW. To the best of our knowledge, there have not been prior reports of anxiogenic effects of *Centella asiatica* in young mice. CAW administered in the diet also had no effect on depressive-like behavior as quantified in the FST.

The limited behavioral effects of CAW in the diet were unexpected. Although we had not previously investigated the effects of CAW on measures of anxiety or depressive like behavior, based on our many published studies on the cognitive effects of CAW a more robust effect at least on those endpoints was expected. We have shown improvements in cognitive function in aged mice following CAW administered in the drinking water at a concentration equivalent to the 200 mg/kg/d dose of CAW used in this study [33, 34] and in a mouse model of Alzheimer’s disease increasing concentrations of CAW in the drinking water (equivalent to the 500 mg/kg/d and 1000 mg/kg/d) resulted in even greater improvements in cognitive function [38]. Interestingly, NORT performance was also improved in both healthy aged mice treated with 200 mg/kg/d equivalent in the drinking water [34] and with all three concentrations administered in the drinking water in the Alzheimer’s disease mouse model [38] whereas that was not seen in the current study where CAW was administered in the diet. This led us to hypothesize that mode of administration could affect behavioral response. We therefore evaluated the effects of the equivalent dose (1000mg/kg/d) of CAW administered in the drinking water to old mice on the same battery of tests of cognitive function, anxiety and depression.

When old mice were treated with 1000 mg/kg/d CAW equivalent in the drinking water, we did observe a significant improvement in both the ODRL and the NORT, and the improvement in ODRL elicited by CAW in the drinking water was even greater than what was seen with CAW (1000 mg/kg/d) in the diet. We also saw significant reductions in anxiety in the open field test but this affect again varied by sex, with aged female mice exhibiting significantly less time in the center than young female mice and aged female mice treated with CAW in the drinking water showing a robust increase in this metric. The age-related increase in time immobile in the open field was attenuated with CAW in the drinking water in both sexes however, this effect is not necessarily related to changes in anxiety exclusively and might instead reflect a change in overall activity with CAW treatment. Intriguingly, CAW in the drinking water significantly increased time in the dark for aged mice in the LD-BOX test suggesting a potential anxiogenic effect. These apparent contradictory effects of CAW in the drinking water on anxiety-like behavior are puzzling. Previous reports of anxiolytic effects of various extracts of the plant have demonstrated improvements in open field and elevated plus maze [21, 22, 25]. LD-BOX was used in a study of a standardized formulation of triterpene compounds (ECa233) from *Centella asiatica* and a reduction in anxiety was observed [20]. However, the concentration of individual triterpenes in ECa233 was much higher (30-50%) than in CAW (< 5%) and the mice tested had been exposed to chronic immobilization stress; both factors may account for the differences in the effect of the CA extracts. Importantly, none of these reports evaluated anxiety in aged animals. In our study age-related increases in anxiety were only detected in the open field and not the LD-BOX. It would be interesting in future studies to employ the elevated plus maze and determine if deficits are evident in aged mice and if CAW in the drinking water attenuates or exacerbates them.

In our study, CAW also had no effect on depressive-like behavior quantified in the FST, regardless of whether it was administered in the diet or in the drinking water. These results are not consistent with the existing literature showing anti-depressant effects of *Centella asiatica* extracts on performance in the FST in a mouse model of olfactory bulbectomy and in healthy young rats [23, 24]. Again, however, these studies were not directly investigating the effects in aged rodents and used other types of CA extracts. It is possible the FST is not sensitive enough to detect age-related changes in depressive-like behavior. If that is the case, then it is unsurprising that a treatment effect would also not be seen in the FST. Future studies could employ other tests of depressive behavior such as the tail suspension, sucrose preference test or conditioned place preference [57], or add an additional stressor to potentially magnify age-related differences, such as restraint stress, cage tilt or noise exposure [58].

One of the most consistent findings of the present study was that CAW effects were more pronounced in the animals that received the extract in the drinking water than those that received it in the diet. A possible explanation for these differences in effect could be differing bioavailability of the constituent compounds of CAW depending on the mode of administration. Further studies are needed to uncover the reason for this differential bioavailability but it is possible that a difference in circulating levels of constituent compounds could explain the marked difference in response to CAW in the drinking water as compared to the diet that we observed in this study.

In summary, these findings suggest a beneficial effect of CAW on age-related changes in cognition and measures of anxiety, but not on depressive-like behavior. The magnitude of these effects was greater when CAW was administered in the drinking water as compared to when it was given in the diet. Future studies are needed to elucidate the mechanism by which CAW elicits its effects on cognition and measures of anxiety as well as to determine if active compounds from the extract reach greater systemic levels when given in the drinking water than in the diet and if these compound levels vary across different brain regions.

## Acknowledgments

This work was funded by NIH-NCCIH grant U19AT010829 and NIH grant S10OD026922. The authors acknowledge Cody Neff and Noah Gladen-Kolarsky for their assistance with the behavioral assays, and Zachary Wiegand of the Oregon State University Department of Food Science and Technology for assistance with the preparation of CAW.

---

**Supplementary Figure 1:**
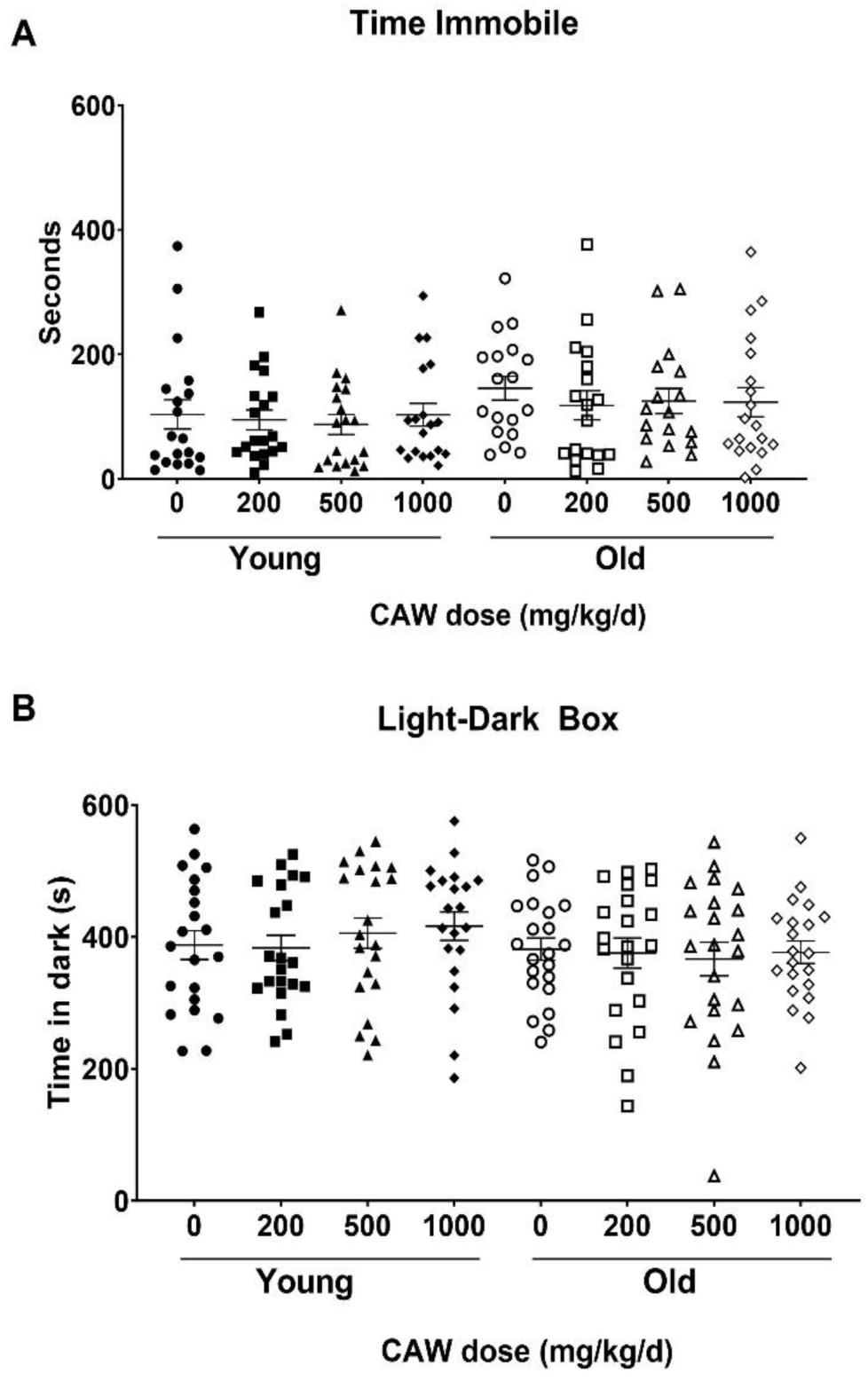
Time immobile in the Open field (OF) test (A) and time in the dark in the Light-Dark Box (LD-Box) were not affected by age or treatment

**Supplementary Table 1:**
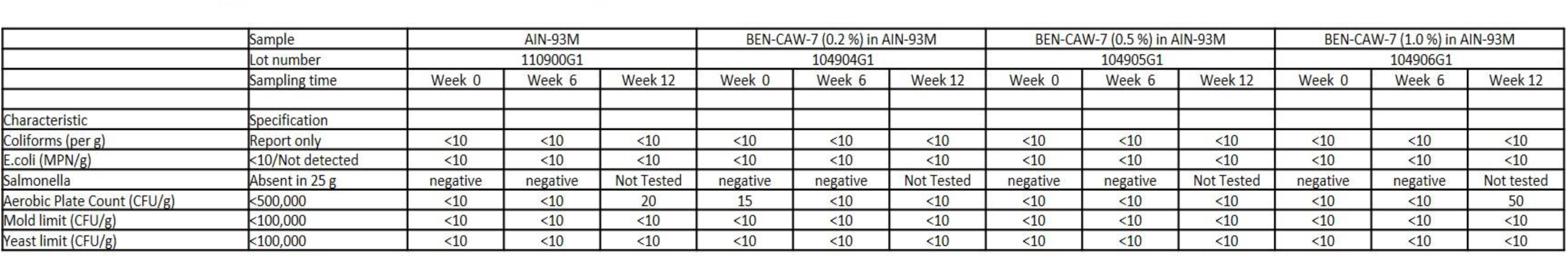
Analysis of Microbial Contamination in Rodent Diet.

**Supplementary Table 2:**
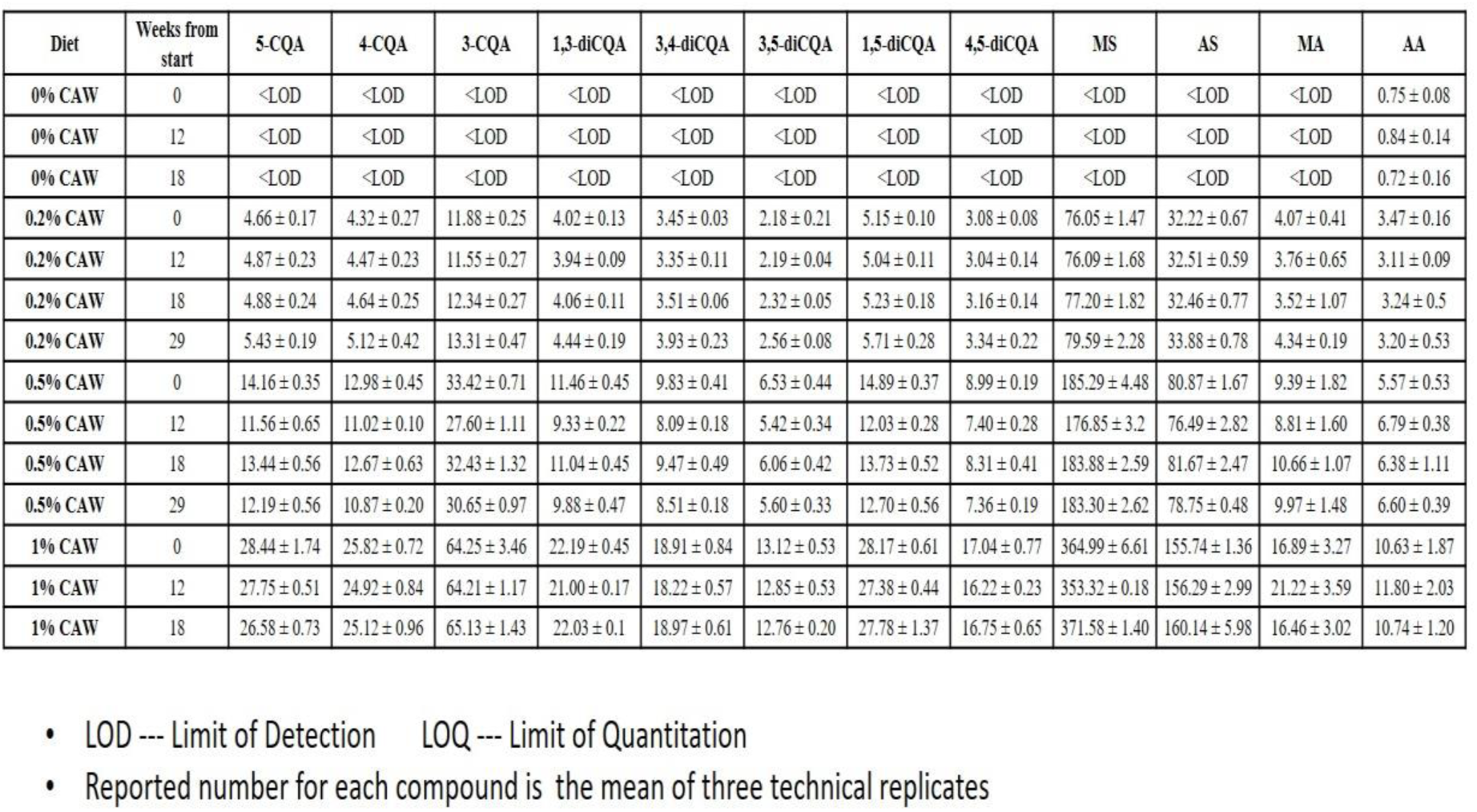
*Centel/a asiatica*marker compounds in Rodent Diet (µg/g)

**Supplementary Table 3:**
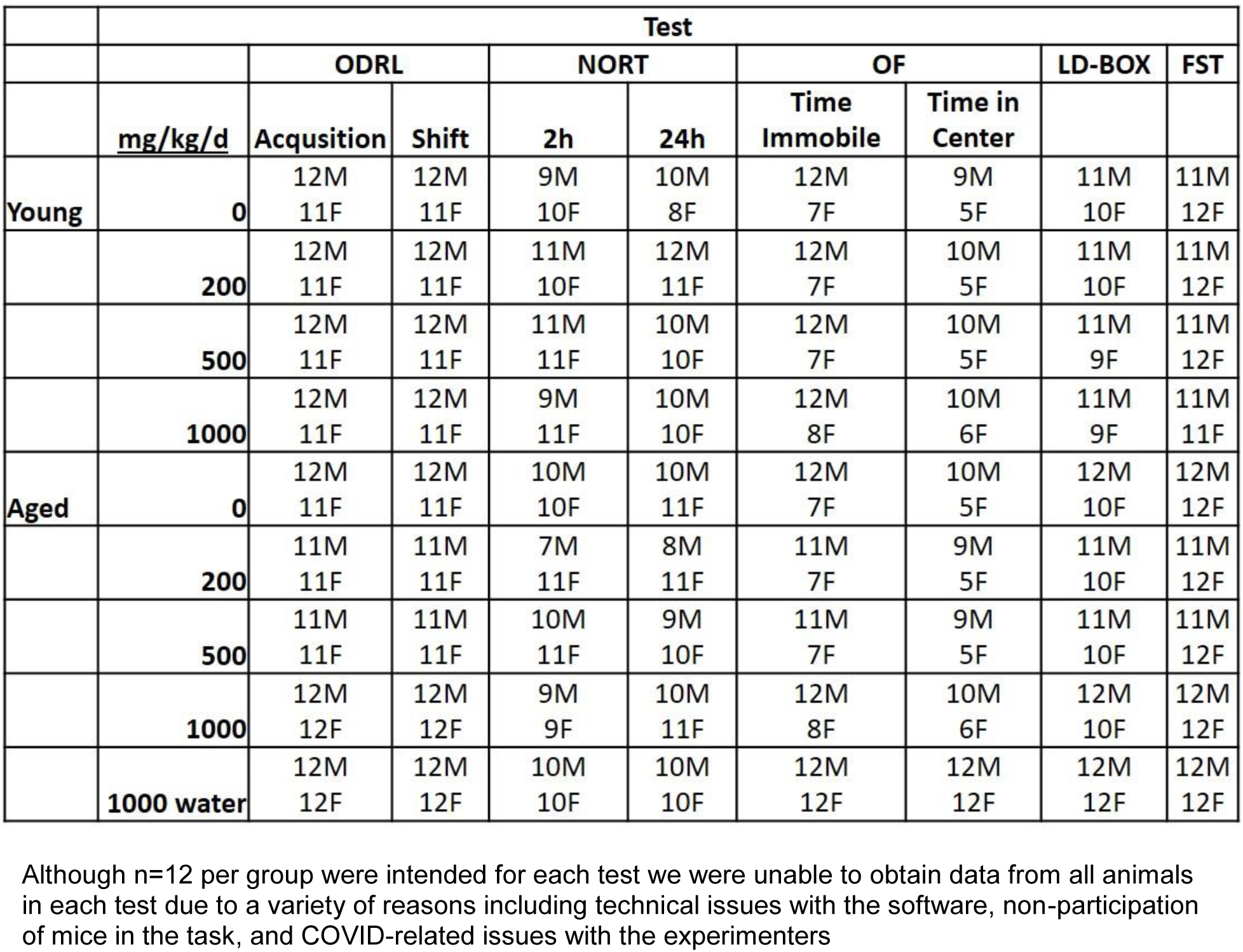
Number of animals that completed each test.

**Supplementary Table 4:**
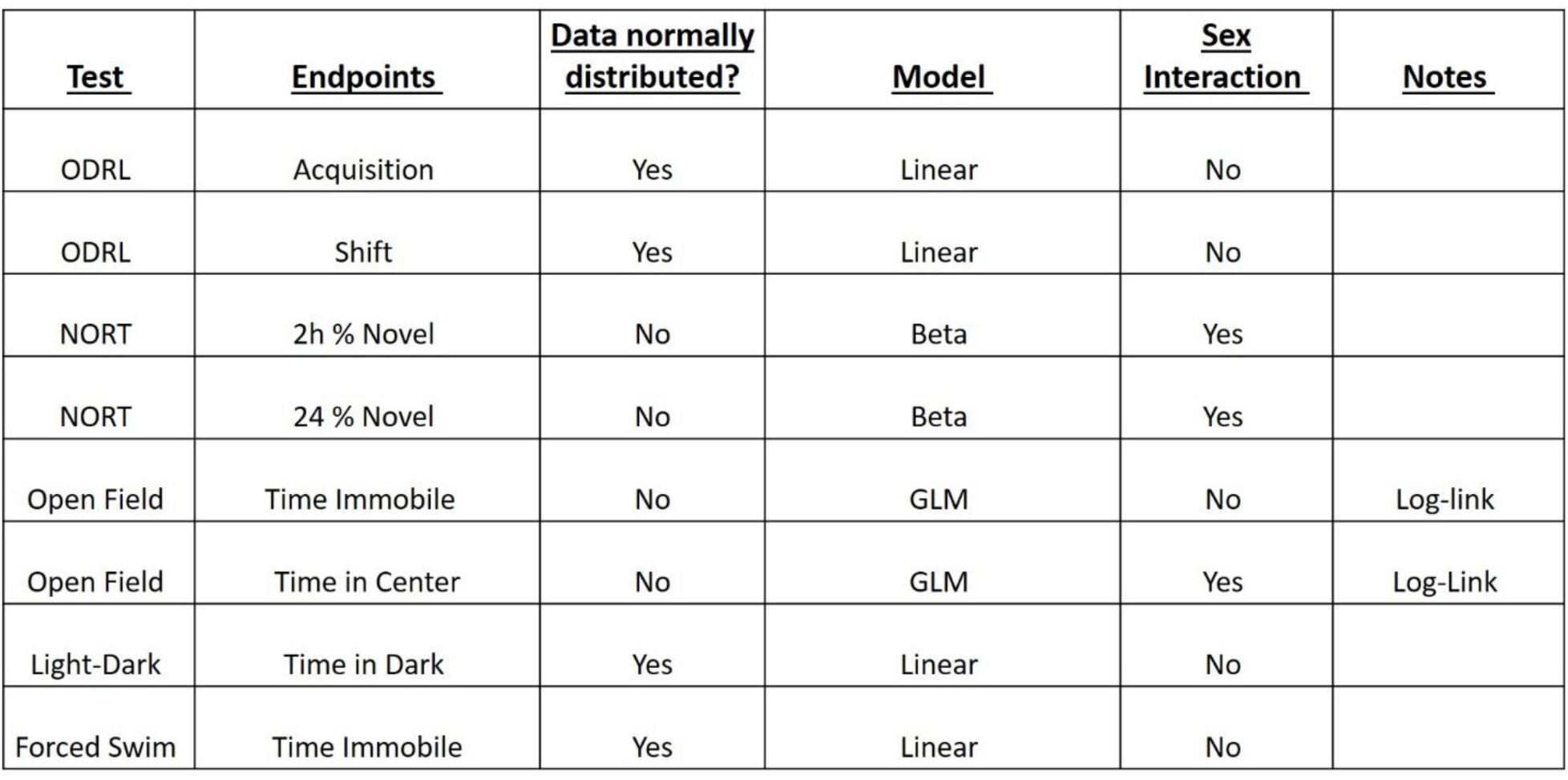
Statistical tests for each endpoint in CAW in the diet dose response experiments:

**Supplementary Table 5:**
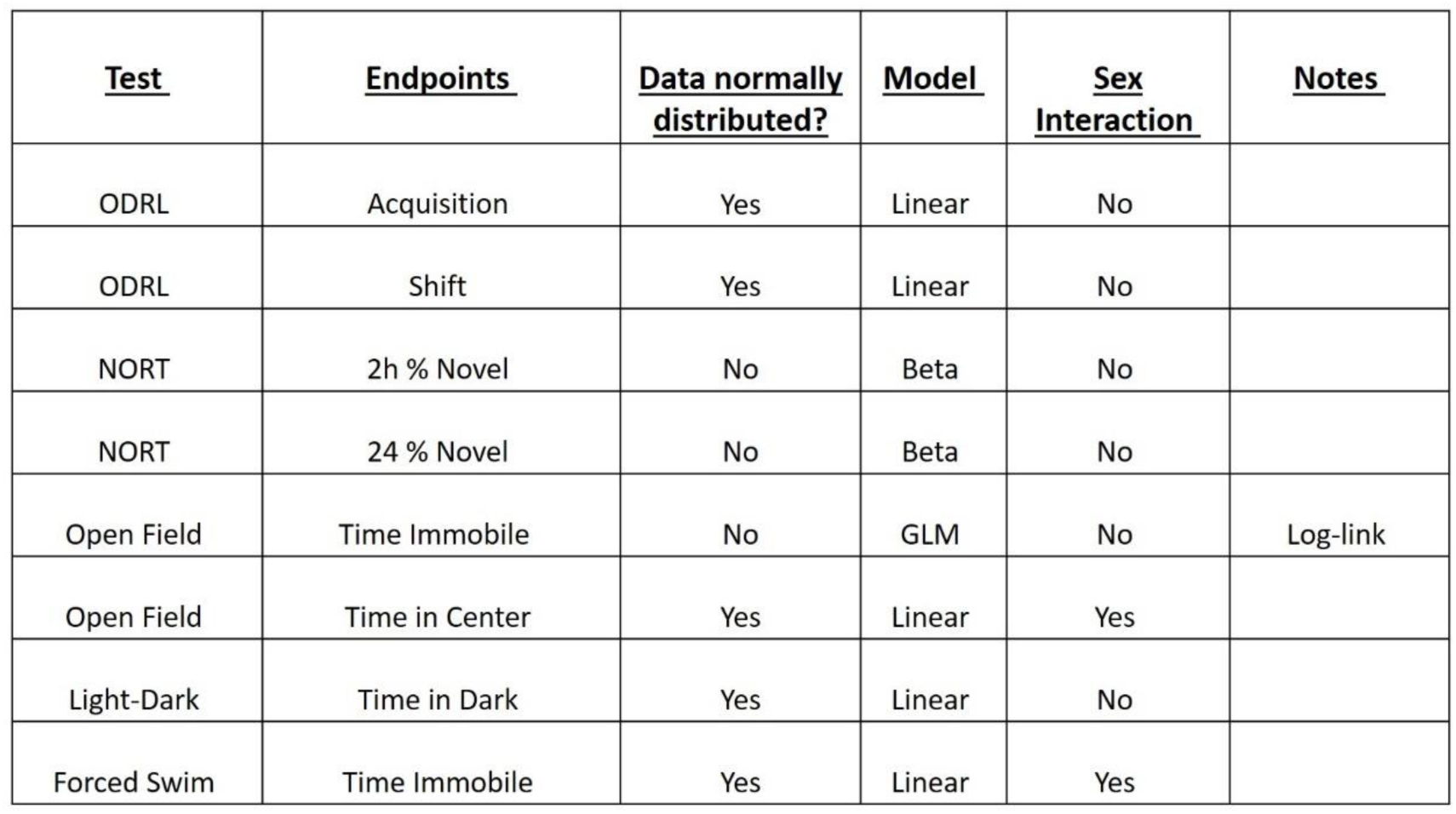
Statistical tests for each endpoint in CAW in the drinking water compared to in the diet experiments:

## Notes

### Competing Interest Statement

Authors Natasha Cerruti and Janis McFerrin are employed by Oregon's Wild Harvest. The remaining authors declare that the research was conducted in the absence of any commercial or financial relationships that could be construed as a potential conflict of interest.

