## Supplementary materials for "Amelioration of age-related cognitive decline and anxiety in mice by *Centella asiatica* extract varies by sex, dose and mode of administration"

### *Supplementary Material*

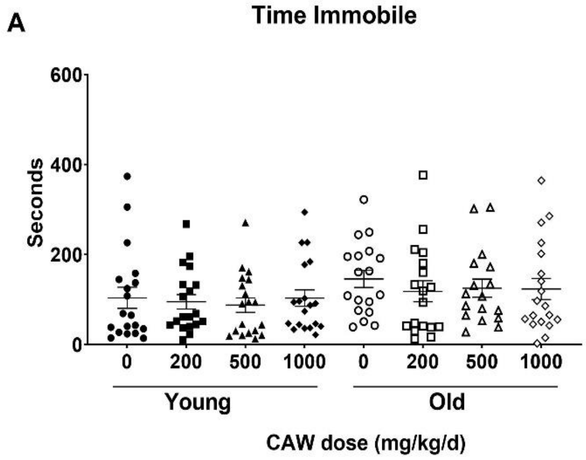

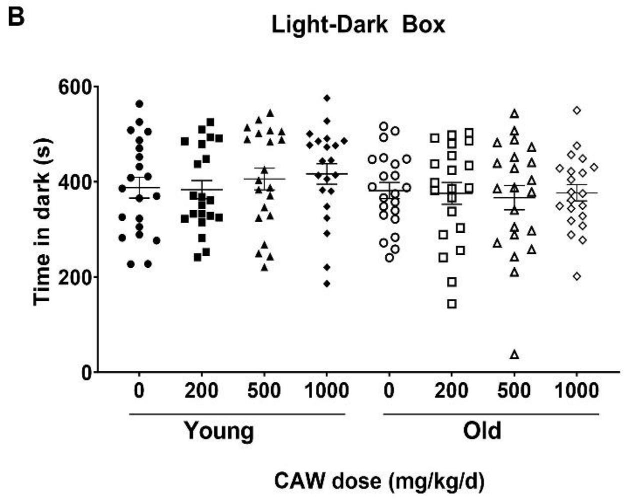

**Supplementary Figure 1: Time immobile in the Open field (OF) test (A) and time in the dark in the Light-Dark Box (LD-Box) were not affected by age or treatment**

#### Supplementary Table 1: Analysis of Microbial Contamination in Rodent Diet

| | Sample | AIN-93M | | | BEN-CAW-7 (0.2 %) in AIN-93M | | | BEN-CAW-7 (0.5 %) in AIN-93M | | | BEN-CAW-7 (1.0 %) in AIN-93M | | |
|---|---|---|---|---|---|---|---|---|---|---|---|---|---|
| | Lot number | 110900G1 | | | 104904G1 | | | 104905G1 | | | 104906G1 | | |
| | Sampling time | Week 0 | Week 6 | Week 12 | Week 0 | Week 6 | Week 12 | Week 0 | Week 6 | Week 12 | Week 0 | Week 6 | Week 12 |
| Characteristic | Specification | | | | | | | | | | | | |
| Coliforms (per g) | Report only | <10 | <10 | <10 | <10 | <10 | <10 | <10 | <10 | <10 | <10 | <10 | <10 |
| E.coli (MPN/g) | <10/Not detected | <10 | <10 | <10 | <10 | <10 | <10 | <10 | <10 | <10 | <10 | <10 | <10 |
| Salmonella | Absent in 25 g | negative | negative | Not Tested | negative | negative | Not Tested | negative | negative | Not Tested | negative | negative | Not tested |
| Aerobic Plate Count (CFU/g) | <500,000 | <10 | <10 | 20 | 15 | <10 | <10 | <10 | <10 | <10 | <10 | <10 | 50 |
| Mold limit (CFU/g) | <100,000 | <10 | <10 | <10 | <10 | <10 | <10 | <10 | <10 | <10 | <10 | <10 | <10 |
| Yeast limit (CFU/g) | <100,000 | <10 | <10 | <10 | <10 | <10 | <10 | <10 | <10 | <10 | <10 | <10 | <10 |

#### Supplementary Table 2: *Centella asiatica* marker compounds in Rodent Diet (µg/g)

| Diet | Weeks from start | 5-CQA | 4-CQA | 3-CQA | 1,3-diCQA | 3,4-diCQA | 3,5-diCQA | 1,5-diCQA | 4,5-diCQA | MS | AS | MA | AA |
|---|---|---|---|---|---|---|---|---|---|---|---|---|---|
| 0% CAW | 0 | <LOD | <LOD | <LOD | <LOD | <LOD | <LOD | <LOD | <LOD | <LOD | <LOD | <LOD | 0.75 ± 0.08 |
| 0% CAW | 12 | <LOD | <LOD | <LOD | <LOD | <LOD | <LOD | <LOD | <LOD | <LOD | <LOD | <LOD | 0.84 ± 0.14 |
| 0% CAW | 18 | <LOD | <LOD | <LOD | <LOD | <LOD | <LOD | <LOD | <LOD | <LOD | <LOD | <LOD | 0.72 ± 0.16 |
| 0.2% CAW | 0 | 4.66 ± 0.17 | 4.32 ± 0.27 | 11.88 ± 0.25 | 4.02 ± 0.13 | 3.45 ± 0.03 | 2.18 ± 0.21 | 5.15 ± 0.10 | 3.08 ± 0.08 | 76.05 ± 1.47 | 32.22 ± 0.67 | 4.07 ± 0.41 | 3.47 ± 0.16 |
| 0.2% CAW | 12 | 4.87 ± 0.23 | 4.47 ± 0.23 | 11.55 ± 0.27 | 3.94 ± 0.09 | 3.35 ± 0.11 | 2.19 ± 0.04 | 5.04 ± 0.11 | 3.04 ± 0.14 | 76.09 ± 1.68 | 32.51 ± 0.59 | 3.76 ± 0.65 | 3.11 ± 0.09 |
| 0.2% CAW | 18 | 4.88 ± 0.24 | 4.64 ± 0.25 | 12.34 ± 0.27 | 4.06 ± 0.11 | 3.51 ± 0.06 | 2.32 ± 0.05 | 5.23 ± 0.18 | 3.16 ± 0.14 | 77.20 ± 1.82 | 32.46 ± 0.77 | 3.52 ± 1.07 | 3.24 ± 0.5 |
| 0.2% CAW | 29 | 5.43 ± 0.19 | 5.12 ± 0.42 | 13.31 ± 0.47 | 4.44 ± 0.19 | 3.93 ± 0.23 | 2.56 ± 0.08 | 5.71 ± 0.28 | 3.34 ± 0.22 | 79.59 ± 2.28 | 33.88 ± 0.78 | 4.34 ± 0.19 | 3.20 ± 0.53 |
| 0.5% CAW | 0 | 14.16 ± 0.35 | 12.98 ± 0.45 | 33.42 ± 0.71 | 11.46 ± 0.45 | 9.83 ± 0.41 | 6.53 ± 0.44 | 14.89 ± 0.37 | 8.99 ± 0.19 | 185.29 ± 4.48 | 80.87 ± 1.67 | 9.39 ± 1.82 | 5.57 ± 0.53 |
| 0.5% CAW | 12 | 11.56 ± 0.65 | 11.02 ± 0.10 | 27.60 ± 1.11 | 9.33 ± 0.22 | 8.09 ± 0.18 | 5.42 ± 0.34 | 12.03 ± 0.28 | 7.40 ± 0.28 | 176.85 ± 3.2 | 76.49 ± 2.82 | 8.81 ± 1.60 | 6.79 ± 0.38 |
| 0.5% CAW | 18 | 13.44 ± 0.56 | 12.67 ± 0.63 | 32.43 ± 1.32 | 11.04 ± 0.45 | 9.47 ± 0.49 | 6.06 ± 0.42 | 13.73 ± 0.52 | 8.31 ± 0.41 | 183.88 ± 2.59 | 81.67 ± 2.47 | 10.66 ± 1.07 | 6.38 ± 1.11 |
| 0.5% CAW | 29 | 12.19 ± 0.56 | 10.87 ± 0.20 | 30.65 ± 0.97 | 9.88 ± 0.47 | 8.51 ± 0.18 | 5.60 ± 0.33 | 12.70 ± 0.56 | 7.36 ± 0.19 | 183.30 ± 2.62 | 78.75 ± 0.48 | 9.97 ± 1.48 | 6.60 ± 0.39 |
| 1% CAW | 0 | 28.44 ± 1.74 | 25.82 ± 0.72 | 64.25 ± 3.46 | 22.19 ± 0.45 | 18.91 ± 0.84 | 13.12 ± 0.53 | 28.17 ± 0.61 | 17.04 ± 0.77 | 364.99 ± 6.61 | 155.74 ± 1.36 | 16.89 ± 3.27 | 10.63 ± 1.87 |
| 1% CAW | 12 | 27.75 ± 0.51 | 24.92 ± 0.84 | 64.21 ± 1.17 | 21.00 ± 0.17 | 18.22 ± 0.57 | 12.85 ± 0.53 | 27.38 ± 0.44 | 16.22 ± 0.23 | 353.32 ± 0.18 | 156.29 ± 2.99 | 21.22 ± 3.59 | 11.80 ± 2.03 |
| 1% CAW | 18 | 26.58 ± 0.73 | 25.12 ± 0.96 | 65.13 ± 1.43 | 22.03 ± 0.1 | 18.97 ± 0.61 | 12.76 ± 0.20 | 27.78 ± 1.37 | 16.75 ± 0.65 | 371.58 ± 1.40 | 160.14 ± 5.98 | 16.46 ± 3.02 | 10.74 ± 1.20 |

- LOD --- Limit of Detection     LOQ --- Limit of Quantitation
- Reported number for each compound is  the mean of three technical replicates

#### Supplementary Table 3: Number of animals that completed each test

| | Test | | | | | | | | |
|---|---|---|---|---|---|---|---|---|---|
| | | ODRL | | NORT | | OF | | LD-BOX | FST |
| | mg/kg/d | Acqusition | Shift | 2h | 24h | Time Immobile | Time in Center | | |
| Young | 0 | 12M 11F | 12M 11F | 9M 10F | 10M 8F | 12M 7F | 9M 5F | 11M 10F | 11M 12F |
| | 200 | 12M 11F | 12M 11F | 11M 10F | 12M 11F | 12M 7F | 10M 5F | 11M 10F | 11M 12F |
| | 500 | 12M 11F | 12M 11F | 11M 11F | 10M 10F | 12M 7F | 10M 5F | 11M 9F | 11M 12F |
| | 1000 | 12M 11F | 12M 11F | 9M 11F | 10M 10F | 12M 8F | 10M 6F | 11M 9F | 11M 11F |
| Aged | 0 | 12M 11F | 12M 11F | 10M 10F | 10M 11F | 12M 7F | 10M 5F | 12M 10F | 12M 12F |
| | 200 | 11M 11F | 11M 11F | 7M 11F | 8M 11F | 11M 7F | 9M 5F | 11M 10F | 11M 12F |
| | 500 | 11M 11F | 11M 11F | 10M 11F | 9M 10F | 11M 7F | 9M 5F | 11M 10F | 11M 12F |
| | 1000 | 12M 12F | 12M 12F | 9M 9F | 10M 11F | 12M 8F | 10M 6F | 12M 10F | 12M 12F |
| | 1000 water | 12M 12F | 12M 12F | 10M 10F | 10M 10F | 12M 12F | 12M 12F | 12M 12F | 12M 12F |

Although n=12 per group were intended for each test we were unable to obtain data from all animals in each test due to a variety of reasons including technical issues with the software, non-participation of mice in the task, and COVID-related issues with the experimenters

**Supplementary Table 4: Statistical tests for each endpoint in CAW in the diet dose response experiments:**

| Test | Endpoints | Data normally distributed? | Model | Sex Interaction | Notes |
|---|---|---|---|---|---|
| ODRL | Acquisition | Yes | Linear | No | |
| ODRL | Shift | Yes | Linear | No | |
| NORT | 2h % Novel | No | Beta | Yes | |
| NORT | 24 % Novel | No | Beta | Yes | |
| Open Field | Time Immobile | No | GLM | No | Log-link |
| Open Field | Time in Center | No | GLM | Yes | Log-Link |
| Light-Dark | Time in Dark | Yes | Linear | No | |
| Forced Swim | Time Immobile | Yes | Linear | No | |

**Supplementary Table 5: Statistical tests for each endpoint in CAW in the drinking water compared to in the diet experiments:**

| Test | Endpoints | Data normally distributed? | Model | Sex Interaction | Notes |
|---|---|---|---|---|---|
| ODRL | Acquisition | Yes | Linear | No | |
| ODRL | Shift | Yes | Linear | No | |
| NORT | 2h % Novel | No | Beta | No | |
| NORT | 24 % Novel | No | Beta | No | |
| Open Field | Time Immobile | No | GLM | No | Log-link |
| Open Field | Time in Center | Yes | Linear | Yes | |
| Light-Dark | Time in Dark | Yes | Linear | No | |
| Forced Swim | Time Immobile | Yes | Linear | Yes | |
